# Denoising broadband CARS hyperspectral images on variance-stabilized data: a unified benchmark

**DOI:** 10.64898/2026.10.09.758096

**Authors:** Haoyu Xu, Marcus T. Cicerone

## Abstract

**Significance:** Broadband coherent anti-Stokes Raman scattering (BCARS) enables label-free hyperspectral chemical imaging, but many signal components are small compared to the accompanying heteroscedastic noise, making optimal denoising crucial.

**Aim:** We seek to identify the highest-performing hyperspectral image denoising methods for BCARS.

**Approach:** We derive a physics-based, variance stabilization transform that yields nearly intensity-independent BCARS noise variance while preserving the dispersive Raman response. Within this framework, we compare six denoisers using a shared phase-retrieval pipeline: singular value decomposition, BM4D, Noise2Noise, deep image prior, self-supervised diffusion, and a cross-covariance-based singular value decomposition (CCV) introduced here. Evaluation uses five simulated cubes with exact ground truth and experimental measurements of glycerol, polystyrene beads, and *Caenorhabditis elegans*.

**Results:** In simulations, the variance stabilization transform reduces the ratio of maximum to minimum channel noise variance from approximately 72 to 1.2. Deep image prior and self-supervised diffusion denoisers achieve the highest mean foreground peak signal-to-noise ratios. CCV matches the mean foreground structural similarity of deep image prior with substantially shorter runtime and no per-cube rank tuning. Experimental comparisons reveal sample-dependent performance.

**Conclusions:** Physics-based variance stabilization provides a common framework for adapting and evaluating hyper-spectral denoisers for BCARS. CCV offers a practical balance of reconstruction fidelity, computational speed, and automated operation.

## 1 Introduction

Raman spectroscopy reads the chemical composition of a sample directly from the vibrational frequencies of its molecular bonds, providing a label-free fingerprint that distinguishes proteins, nucleic acids, and lipids without stains or genetic modification.^1, 2^ Its spontaneous cross-section is tiny because the field for one of the two transitions involved is supplied by vacuum fluctuations.^3^ Thus, acquiring a full spectrum at every image pixel is slow, and this weakness has long confined spontaneous Raman microscopy to low throughput. Coherent Raman scattering removes this limit by providing strong laser fields for both transitions, thus amplifying the vibrational signal by orders of magnitude and enabling chemically specific imaging at biologically relevant speeds.^4, 5^ Two modalities dominate, stimulated Raman scattering (SRS) and coherent anti-Stokes Raman scattering (CARS). The SRS signal is discriminated from a large, unconverted laser background through pump modulation, and the modulated, vibrationally resonant SRS signal is linear in concentration.^6–8^ CARS is a parametric process that carries a smaller, nonresonant background (NRB) that is difficult to remove experimentally.^9, 10^ It leaves a purely resonant signal that is quadratic in concentration, whereas phase-retrieval methods are used to obtain a resonant susceptibility that is linear in concentration.^11, 12^

Broadband CARS (BCARS) exploits the parametric nature of the CARS process to generate and detect vibrational coherence over a wide spectral range. It uses a ∼15 fs laser pulse to impulsively excite vibrational modes in the fingerprint spectral region (below 1800 cm*^−^*^1^), and excites the “CH stretch” spectral region (2500–3200 cm*^−^*^1^) through the interaction of the ∼15 fs and a blue-shifted, spectrally narrow, ∼3 ps pulse. The ps pulse is then reused as a probe for the entire coherence, leading to the generation of anti-Stokes signal with sufficient breadth to record a complete vibrational spectrum spanning thousands of wavenumbers at every spatial pixel in a single acquisition^13–16^ (Fig. 1). This approach produces hyperspectral data cubes that resolve the full chemical fingerprint and CH-stretch region of single cells, intact tissues, and whole organisms such as *Caenorhabditis elegans*.^15, 17–19^

**Fig 1.**
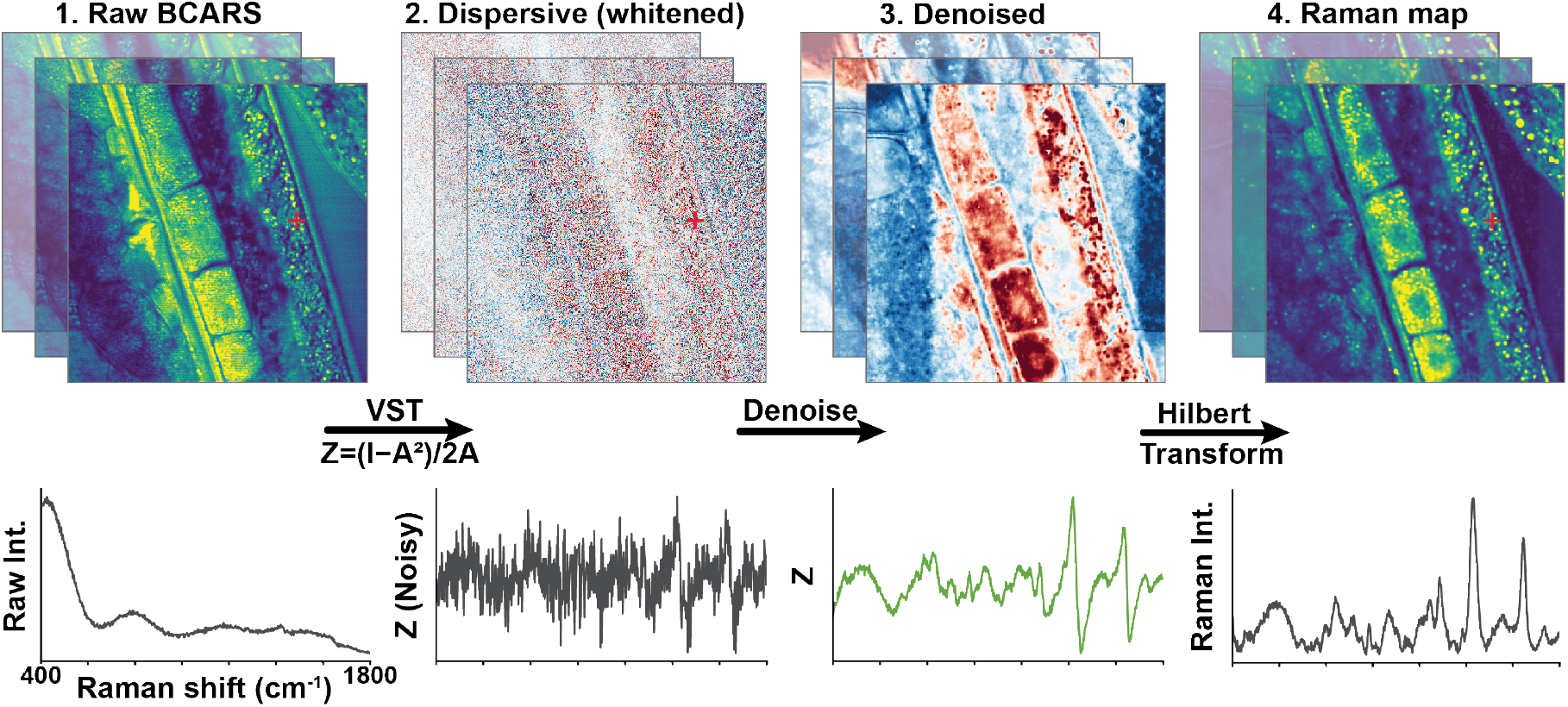
BCARS denoising benchmark and shared processing pipeline. Raw BCARS hyperspectral acquisitions (1), dominated by the nonresonant background, are converted to the signed dispersive cube 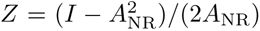 by the variance-stabilizing transform (2), which equalizes the channel-dependent noise while placing the signal in units of Re *χ*_R_. Six denoisers are applied in (3), after which a Hilbert transform and phase-error correction recover the Raman-like resonant map (4). Bottom row: a representative single-pixel spectrum at each stage (raw intensity; noisy dispersive *Z*; denoised *Z*; retrieved Raman intensity), with the red marker indicating the sampled pixel. Every method enters at stage 3 and is scored after stage 4, so that, apart from the bounded-range normalization DDS2M requires (Sec. 3.2), the denoiser is the only variable in the comparison.

This richness comes at a cost. The fingerprint region of the BCARS spectrum carries the sharp, chemically specific bands that separate molecular species and molecular conformation states such as protein secondary structure,^20^ nucleic acids,^21^ and lipid ordering^22^ that the broad CH-stretch region near 2800–3200 cm*^−^*^1^ cannot resolve.^17, 23^ However, these fingerprint Raman signals are typically an order of magnitude smaller than the CH-stretch signals and can be two orders of magnitude weaker than the nonresonant background that they are superimposed on. The impulsive excitation scheme we use^24, 25^ creates strong coherence across the fingerprint window (Fig. S1), efficiently exciting these vibrational modes, but also generating a strong NRB. While the vibrational signal is heterodyne-amplified by the NRB, its inherently low signal-to-noise is not improved. Thus, these fingerprint bands are the weakest and have the highest noise content in the entire BCARS signal.^5, 15^ At the millisecond dwell times required for high-throughput imaging, recovering quantitative chemical information from this spectral region hinges on reliable and effective denoising, which is what this work addresses.

Two coupled physical obstacles hinder the extraction of quantitative vibrational spectra from raw BCARS cubes. First, as noted above, the resonant third-order nonlinear susceptibility (*χ*_R_) carrying the quantitative molecular vibrational fingerprint is phase-mixed with, and typically dwarfed by, a nonresonant background (*χ*_NR_) that originates from electronic four-wave mixing of the host medium.^5, 16^ Because the detected signal scales with |*χ*_R_ + *χ*_NR_|^2^, the weak vibrational features appear as small dispersive modulations riding on top of a much larger baseline. Second, BCARS spectra are contaminated by both photon shot noise and detector readout noise, yielding a heteroscedastic Poisson–Gaussian noise structure where the local noise variance increases with the signal intensity.^26^

To reduce noise in coherent Raman hyperspectral data, low-rank matrix approximations such as truncated singular value decomposition (SVD) have long served as the standard workhorse, exploiting the high correlation in spatial amplitude patterns across continuous spectral channels.^27^ Truncated SVD minimizes the squared reconstruction error across the data cube. With heteroscedastic noise, high-variance channels can dominate this objective and obscure weaker fingerprint features. Variance stabilization, commonly performed with an Anscombe transform,^28, 29^ reduces this imbalance before SVD is applied. Because the transform is nonlinear, the denoised data must then be returned to intensity units with a bias-corrected inverse.^29^ A practical complication of SVD on fingerprint bioRaman spectra is that the signal and noise levels are similar for many peaks, choosing singular vectors to include as much signal from those weak peaks as possible is often an operator-tuned choice. Because the Anscombe transform only approximately stabilizes heteroscedastic noise, residual noise variance can be redistributed into subspaces that overlap with weak spectral features, complicating rank selection and potentially reducing recovery of low-amplitude peaks.

Recently, deep learning models including Noise2Noise (N2N),^30^ deep image prior (DIP),^31, 32^ and diffusion-based generative models^33, 34^ have emerged for general hyperspectral restoration. In coherent Raman imaging, deep-learning denoising has been demonstrated in CARS endoscopy^35^ and in hyperspectral SRS, using supervised spatial–spectral learning,^36^ unsupervised single-image learning,^37^ and Noise2Noise;^38^ to our knowledge, diffusion-based denoisers have not been applied to coherent Raman data, and none of these approaches has been compared on BCARS.

Here, we use a set of synthetic and experimental BCARS data to evaluate the performance of the computational denoisers mentioned above and another we introduce, cross-covariance SVD (CCV), which builds on inter-battery factor analysis.^39^ Putting these on a common footing requires equalizing the underlying noise variance prior to spectral restoration (Fig. 1). To do this, we introduce a variance-stabilizing transform (VST) derived from the BCARS forward model that linearizes detected intensity about the NRB-dominant limit and, in the same step, maps the intensity-dependent Poisson–Gaussian noise to approximately homoscedastic noise, which is close to Gaussian at the photon counts of our measurements. Given a measured NRB, the transform inverts exactly. By performing denoising entirely inside the VST-whitened dispersive domain, linear subspace algorithms (SVD, CCV), patch-based collaborative filtering (block-matching and 4-D filtering, BM4D),^40^ self-supervised convolutional networks (N2N), untrained convolutional priors (DIP), and self-supervised diffusion models (denoising diffusion spatio-spectral model, DDS2M) are evaluated on an identical statistical footing where errors are weighted equally across bright and dark channels. Furthermore, every denoised estimate is subsequently passed through a single, shared phase-retrieval pipeline, so that measured differences in spectral and spatial fidelity are attributable to the restoration algorithm itself.

We apply this benchmark to five data cubes that together provide exact ground truth and realism: five simulated cubes with exact complex ground truth (based on an experimental BCARS image), homogeneous glycerol, polystyrene beads of known size, and *C. elegans* tissue. Performance depends on the sample, evaluation metric, and image region. DDS2M and DIP achieve the highest mean foreground PSNR on the simulated cubes, while CCV combines competitive foreground fidelity with automatic component selection and shorter runtime. The experimental comparisons also identify sample-dependent artifacts that are not captured by aggregate simulation metrics. Figure 6 summarizes these trade-offs in fidelity, runtime, and operator input.

## 2 Theory

Broadband coherent anti-Stokes Raman scattering (BCARS) encodes the vibrational spectrum in a signed, background-dominated interference term, and this study compares the performance of each denoiser in acting on that term. At spectral channel *i* the detector records an intensity *I_i_* ∝ |*χ_i_*|^2^ proportional to the third-order susceptibility *χ_i_* = *χ*_NR_*_,i_* + *χ*_R_*_,i_*, where the nonresonant background *χ*_NR_*_,i_* ∈ ℝ*_>_*_0_ is real and the resonant part *χ*_R_*_,i_* ∈ ℂ carries the Raman response.^15, 17^ Expanding the modulus,

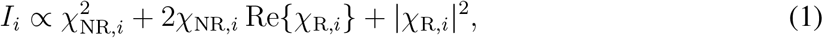

separates the measurement into a nonresonant background 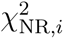, a resonant–nonresonant interference 2*χ*_NR_*_,i_* Re{*χ*_R_*_,i_*} that is linear in Re{*χ*_R_*_,i_*}, and a homodyne term |*χ*_R_*_,i_*|^2^. In the small-signal regime relevant to bioRaman fingerprint spectra, |*χ*_R_*_,i_*| ≪ *χ*_NR_*_,i_*: the background dominates the intensity, the homodyne term is negligible, and the Raman information is extracted from a small signed dispersive signal riding on a large pedestal.

The amplitude and noise of this background pedestal are channel-dependent. The detected signal follows a Poisson–Gaussian model: the recorded value *Y_i_*is a scaled photoelectron count with Poisson-distributed shot noise plus signal-independent Gaussian-distributed readout noise, so that its mean is *µ_i_*= *I_i_* and its variance is

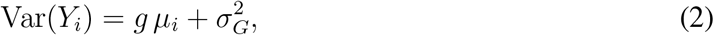

where *g* is the effective gain of the recorded data (the product of the detector conversion and any global intensity normalization) and 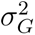 is the readout variance in the same units. Writing the mean as *µ_i_* = *µ*_0_*_,i_* + *δ_i_*, with nonresonant baseline 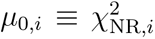 and resonant perturbation *δ_i_* = 2*χ*_NR_*_,i_* Re{*χ*_R_*_,i_*} + |*χ*_R_*_,i_*|^2^, the noise is set almost entirely by the baseline, 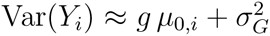 the shot-noise standard deviation scales with the square root of the baseline amplitude, 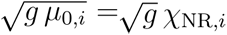 Because the spectral profile of the excitation varies by nearly two orders of magnitude across the window, the noise is strongly heteroscedastic: bright channels carry far larger absolute noise than channels with weak signal.^15^ Applied to the raw cube, any low-rank or covariance-based denoiser would therefore be steered by the brightest channels rather than by signal-to-noise content, and this bias must be removed before a fair comparison across methods is possible.

We remove the pedestal and equalize the noise with a single variance-stabilization transform (VST), applied pixel by pixel:

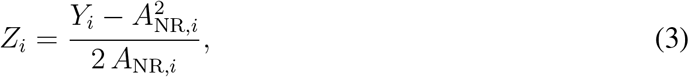

with 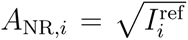 the nonresonant amplitude obtained from a reference cube acquired on water, used as an approximately nonresonant reference. Estimating the pedestal from an independent measurement rather than from the sample itself guarantees that the baseline cannot absorb sample resonances; the residual mismatch between the reference and the sample background is slowly varying and is removed downstream by the shared front-end (Section 3.2). The numerator subtracts the background pedestal; the denominator does two jobs at once. First, in Eq. (1) the interference term amplifies the resonant signal by the factor 2*χ*_NR_*_,i_*; normalizing by this factor expresses *Z* directly in units of Re{*χ*_R_}, neglecting the homodyne term, the signal-bearing part of *Z_i_* is Re{*χ*_R_*_,i_*}. Second, because the shot-noise variance is proportional to the same amplitude, the division equalizes the noise across channels. Propagating Eq. (2) through the transform and using 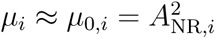,

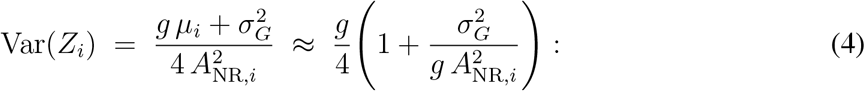

wherever shot noise dominates 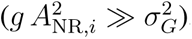, the variance sits at the constant plateau *g/*4, independent of the channel. The gain sets a global scale that is accommodated by the denoisers’ scaling properties or input normalization. The transformed cube has approximately channel-independent noise variance and a signal that is linear in Re{*χ*_R_} to first order. Subsequent baseline subtraction and normalization produce the denoiser inputs described in Section 3.2.

We retain the term variance-stabilizing transform because Eq. (3) serves the same purpose as the pointwise nonlinear transforms conventionally used for Poisson–Gaussian data, such as the generalized Anscombe transform:^26^ it maps a signal-dependent noise variance onto an approximately constant one. It differs in construction. Because the channel dependence of the noise is set here by a separately measured nonresonant amplitude rather than by the unknown signal, stabilization is achieved by an affine rescaling with a known, deterministic factor. Two practical consequences follow. The transform inverts exactly, with none of the bias correction that Anscombe-type inverses require, and it leaves the signal linear in Re{*χ*_R_} to first order.

Fig. 2 validates the transform against its own noise model. Before whitening, the per-channel variance traces the nonresonant envelope predicted by Eq. (2), spanning 1.17×10*^−^*^5^ to 8.43×10*^−^*^4^, a factor of 72 across the window (A), with the empirical and theoretical curves in close agreement and visibly different noise widths at high-, mid-, and low-wavenumber channels (*σ*^2^ = 8.1 × 10*^−^*^5^, 1.8 × 10*^−^*^4^, 3.7 × 10*^−^*^4^; C). After whitening the same range narrows to 2.02 × 10*^−^*^4^–2.37 × 10*^−^*^4^, a factor of 1.2 (B), and the three distributions collapse onto a common width (C); the plateau agrees with the value *g/*4 = 2.04 × 10*^−^*^4^ predicted by Eq. (4). The transform therefore reduces the channel-to-channel variance spread by a factor of roughly 60 and lands on the theoretically expected level. These panels use the simulated cube, where the true per-channel mean *µ_i_* is known and the noise model can be checked exactly; the same stabilization holds on measured data, where the per-channel variance of an experimental worm cube, normalized to its mean across wavenumbers, falls from a roughly 6× spread to the ideal unit line (D). The stabilized variance rises toward 1700–1800 cm*^−^*^1^, the Raman-silent edge of the window where the nonresonant amplitude *A*_NR_ is smallest. The read-noise term 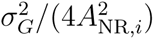 in Eq. (4) becomes appreciable as the background amplitude decreases. Agreement between the theoretical and empirical curves supports read noise as the explanation for this rise.

**Fig 2.**
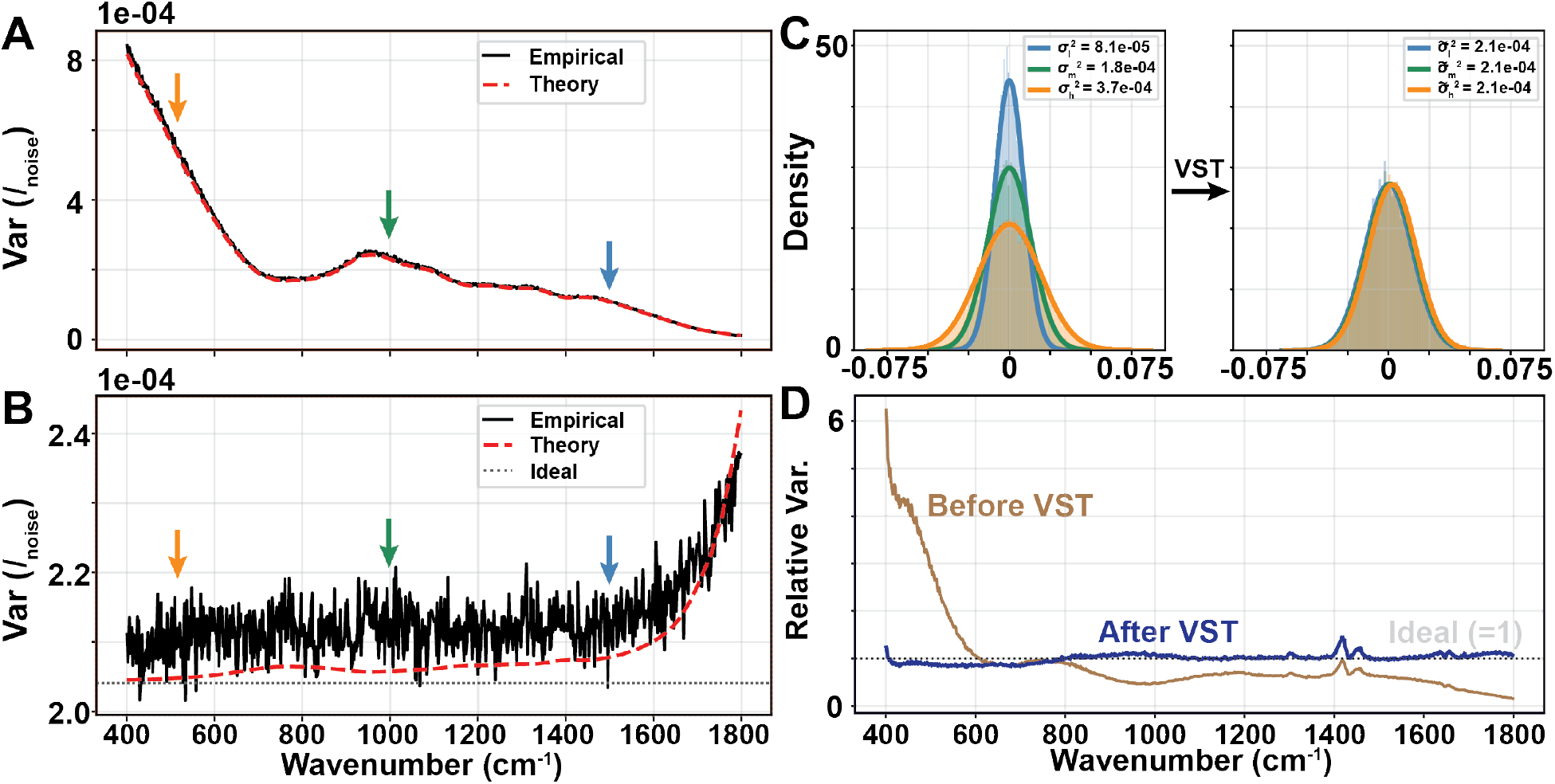
Variance-stabilizing whitening of the BCARS noise. Per-channel noise variance versus wavenumber before (A) and after (B) the VST on the simulated worm cube, with the empirical variance (black), the Poisson–Gaussian model of Eq. (2) (red, “theory”), and the constant-variance plateau of Eq. (4) (dotted). Arrows mark the high-, mid-, and low-wavenumber channels sampled at right. The stabilized variance is flat except toward 1700–1800 cm*^−^*^1^, where the nonresonant amplitude *A*_NR_ is smallest and the readout term of Eq. (4) becomes appreciable. (C) Noise distributions at the three marked channels, heteroscedastic before the VST (*σ*^2^ = 8.1 × 10*^−^*^5^, 1.8 × 10*^−^*^4^, 3.7 × 10*^−^*^4^) and collapsed to a common width after 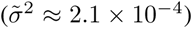. (D) Relative variance across the band flattens from ∼ 6× to the ideal unit line in experimental data.

The stabilization is therefore only locally homoscedastic, and its accuracy rests on *δ_i_/µ*_0_*_,i_* being small. A practical criterion is |*χ*_R_*_,i_*|*/χ*_NR_*_,i_* ≲ 0.1, corresponding, to first order, to at most a 20% perturbation of the stabilized variance; the approximation degrades at strong resonances and at the weak-background spectral edges.

## 3 Methods

### 3.1 Datasets

We evaluate the denoisers on four BCARS hyperspectral datasets, chosen so that the comparison spans a controlled setting with exact ground truth and progressively more realistic samples without it. All cubes cover the fingerprint region from 400 to 1800 cm*^−^*^1^ in roughly 690 spectral channels, at a spectral resolution of 6–7 cm*^−^*^1^. The experimental cubes were acquired on the BCARS microscope described previously,^24, 25^ with a pixel size of 0.26 µm and a dwell time of 4 ms per pixel, using 30 mW of probe and 50 mW of supercontinuum at the sample. A nonresonant reference cube is acquired on water under the same conditions and supplies the amplitude used by the variance-stabilizing front-end (Section 3.2).

The simulated data are the only dataset with an exact ground truth. They are built physically: model Raman spectra obtained by peak fitting^41^ are combined with a known NRB to form noise-free complex CARS spectra, to which Poisson and Gaussian noise are added to reproduce the measured noise statistics. We generate five such cubes, corresponding to five distinct BCARS images, and report every simulated-data metric as a mean and standard deviation across them. The NRB used in the forward model is the same one supplied to the variance-stabilizing front-end, so the reference mismatch that affects experimental cubes is absent here by construction, and the noise model of Section 2 can be checked exactly. Because the underlying susceptibility is known exactly, these cubes furnish a true complex ground truth; we score the denoisers on the resonant (Im{*χ*_R_}) lineshape, the quantity from which the Raman signal is ultimately read (Section 3.5).

The three experimental cubes trade exact ground truth for realism. The spatial homogeneity of glycerol allows us to construct a measurement-based pseudo-ground-truth by averaging the noisy spectra along the slow scan axis. The polystyrene-bead cube contributes sharp, high-contrast structure of known size, the beads having a nominal diameter of 1 µm, and the bead edges provide a test of spatial fidelity and artifact generation. The experimental *C. elegans* cube provides heterogeneous real tissue for qualitative comparison of background fluctuations, tissue morphology and fine spatial detail after denoising. Five further *C. elegans* cubes, acquired under the same conditions, are used only to train the N2N network (Section 3.3).

### 3.2 Shared Preprocessing

All denoisers use the same variance stabilization, baseline subtraction, and normalization procedure. DDS2M additionally requires clipping to its supported input range, as described below. Raw BCARS acquisitions are converted to a quantitative dispersive quantity following the standard analysis chain for the technique,^11, 15^ with a variance-stabilizing modification detailed in Section 2; we summarize the three stages the front-end applies. First, a per-pixel variance-stabilizing transform, divided by the amplitude of a nonresonant reference cube measured on water, maps the heteroscedastic Poisson–Gaussian measurement to the signed dispersive susceptibility Re *χ*_R_ with approximately constant, channel-independent (homoscedastic) noise (Fig. 2), so that the linear low-rank methods and the modern learned methods alike act on nearly Gaussian noise. Second, a wide symmetric Whittaker baseline subtraction,^42^ with the smoothing parameter held fixed across all four cubes, removes the low-frequency background left by the mismatch between the NRB reference and the BCARS baseline. After such baseline subtraction, the dispersive bipolar lineshape is preserved (Fig. 1 panel 2). Third, a robust-symmetric normalization maps the signed cube to [0, 1] with zero mapped to 0.5, scaling the positive and negative lobes identically by the 0.5th and 99.5th percentiles of the cube. We keep two variants of this normalization that differ only in the bright tails: a clip cube hard-limited to [0, 1], and a noclip cube that preserves peak amplitudes reaching several times the nominal maximum; the distinction matters only for methods whose output range is intrinsically bounded: DDS2M clamps its output to [0, 1] and is therefore run on the clip cube (Section 5). The resulting cube is the common input to the denoisers, and the full pipeline is summarized in Fig. 1.

### 3.3 Denoisers under comparison

The benchmark covers six denoisers of increasing modeling capacity, from a linear low-rank baseline to self-supervised deep networks, drawn from five distinct denoising families. All operate on the same whitened dispersive cube (Section 3.2); the network architectures, hyperparameters, and any deviation from the reference implementations of the learned methods are collected in Table S1. All runtimes quoted in Section 5 were measured on a single NVIDIA RTX 5090 GPU with an Intel Core i9-12900K CPU and 64 GB of RAM. We summarize each method below and defer the operator-free rank-selection rule we contribute to Section 3.4.

Truncated singular value decomposition (SVD) is the standard multivariate denoiser for BCARS and serves as our baseline. The whitened cube is flattened to a pixel-by-wavenumber matrix and approximated by its leading singular components, which capture the few chemical spectra shared across many pixels while the noise is spread over the discarded components.^15, 27^ In our SVD baseline, we manually select the visually signal-bearing singular components; CCV provides an alternative based on split-half reproducibility (Section 3.4).

BM4D brings in the spatial self-similarity that a global low-rank model ignores. Representing the nonlocal, patch-based denoising family, it groups similar three-dimensional patches from across the cube and applies a collaborative transform-domain shrinkage.^40^ The method assumes additive white Gaussian noise, an assumption the whitening front-end is designed to satisfy; we set its single noise-level parameter from the standard deviation measured in a signal-free spatial region of the whitened cube.

Noise2Noise (N2N) is a self-supervised convolutional network that learns to map one noisy realization of a scene to a second, statistically independent realization of the same scene; trained with a mean absolute error (MAE) loss (Table S1), the network converges to the per-pixel conditional median which is also the clean signal for the approximately symmetric, zero-mean noise after the front-end without seeing clean targets.^30^ The same principle has been applied to hyperspectral coherent Raman imaging by the SPEND method,^38^ which constructs the two required views by an odd/even split of the spectral channels. We adopt this construction and therefore include N2N as a reproduction and baseline: we form the two views by the same odd/even split of the whitened cube and train a three-dimensional U-Net^43^ (Table S1). N2N is the only method in the benchmark trained on a corpus separate from the cube it restores; the experimental model is trained on five *C. elegans* cubes acquired under the same conditions and disjoint from the benchmark cubes. To test whether its behavior on sharp features depends on the training distribution, we train a second model with the 1 µm bead cube added to that corpus and compare the two on the bead data (Fig. S5).

The deep image prior (DIP) fits an untrained convolutional network to a single noisy cube, relying on the network’s bias toward spatially coherent structure to learn the signal before it fits the noise, so that early stopping yields a denoised estimate.^31^ Following S2DIP,^32^ we regularize the prior with a total-variation term, restricted here to the wavenumber axis to enforce spectral smoothness (Table S1). The iteration count is chosen by visual inspection of the reconstructed cube; the absence of a ground-truth-free stopping rule is taken up in Section 5. To our knowledge, this is the first application of DIP to BCARS denoising.

DDS2M is a self-supervised diffusion model for hyperspectral restoration that couples an untrained spatial network with an untrained spectral network and denoises through a reverse diffusion trajectory.^33^ It requires no external training data.^33^ Like DIP, it has not previously been applied to BCARS; its computational cost and its bounded output range are examined in Section 5.

### 3.4 Cross-covariance singular value decomposition (CCV)

We introduce cross-covariance-based SVD (CCV), which constructs a low-rank representation from two spectral views of a single acquisition. Under the independent-noise assumption, the noise–noise contribution to their cross-covariance vanishes in expectation. Agreement between the corresponding score images provides a dimensionless criterion for component selection.

We construct two spectral views from a single acquisition. The BCARS spectral resolution is 6–7 cm*^−^*^1^, whereas the camera samples the spectrum at roughly 2 cm*^−^*^1^ per channel. This oversampling provides adjacent channels with similar signal content. We additionally assume that their noise is independent. We note that spectrally correlated noise, e.g., from detector cross-talk would violate this assumption, but we find no evidence for such a violation. As with N2N, we form the two spectral image views by an odd/even spectral channel split (Section 3.3): an even-channel block *X_E_* and an odd-channel block *X_O_*, each in ℝ*^N×m^* with *m* = ⌊*M/*2⌋ for a cube of *N* pixels and *M* channels. After column-centering each block (*X̃_E_, X̃_O_*), we take the cross-covariance and its singular value decomposition,

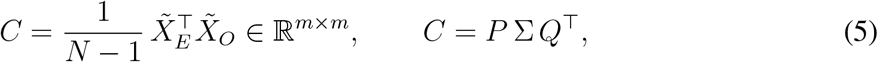

and project each view onto its modes to obtain per-pixel score images 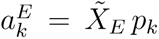 and 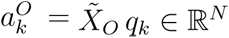.

In conventional SVD, the decomposition is based on the autocovariance matrix *X^⊤^X*, which contains the noise term *E^⊤^E*. Because E[*E^⊤^E*] ≠ 0, noise introduces a full-rank bias that inflates all singular values and mixes signal and noise across the singular-vector basis.

In contrast, CCV is performed on the cross-covariance constructed from two nearly redundant spectral representations with independent noise.

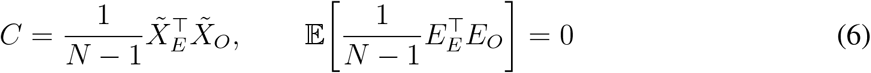

The noise autocovariance term is eliminated, leaving only zero-mean fluctuations that decrease as *N ^−^*^1^*^/^*^2^. Consequently, CCV produces singular values with substantially reduced noise bias and a noise floor that decreases approximately as 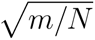, leading to improved separation of signal and noise. Signal modes remain correlated between the two spectral copies, whereas noise modes are only weakly correlated, providing a criterion for rank selection.

Truncated SVD denoises the whitened cube but leaves open how many components to keep: on real data the singular-value spectrum may have no clean break that marks the signal/noise boundary, so the rank is set by the operator.^15, 27^ Automatic alternatives exist: optimal singular-value hard thresholding under a Marchenko–Pastur noise model,^44^ Malinowski’s indicator function,^45^ and parallel analysis^46^ all select a rank without operator input, but each requires either a noise-level estimate or an assumed null spectrum against which the singular values are compared.

We replace the operator’s choice with an objective, scale-free criterion that requires no noise-level estimate, which we term cross-covariance rank selection. The criterion rests on reproducibility of signal across two views of the same measurement that carry independent noise.

A component is kept when its two score images agree, measured by their split-half correlation

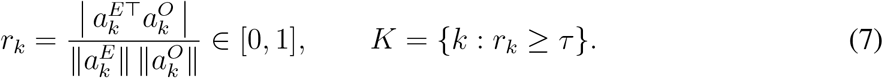

For a mode with per-component signal-to-noise ratio SNR*_k_*, the split-half correlation approximately follows the classical reliability relation *r_k_* = SNR*_k/_*(SNR*_k_* + 1);^47^ we use *τ* = 0.5 (SNR*_k_* ≈ 1) as an empirical selection rule. Because the modes are fitted on the same data, *r_k_*is biased upward for noise modes when *N* is not much larger than *m*; for our cubes *N* ≫ *m*. Unlike an absolute singular-value cutoff, the correlation threshold is dimensionless; we use *τ* = 0.5 throughout. On pre-whitened data, whose noise is approximately isotropic across channels, *r_k_* is the split-half reliability of the cross-covariance modes, and the rule *r_k_* ≥ *τ* retains exactly those modes that reproduce across the two views.

Given the retained set *K*, rebuilding the two halves independently and interleaving them back would drive adjacent channels with two different coefficients (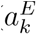 versus 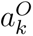); the parity-dependent mismatch oscillates at the spectral Nyquist frequency as a period-2 comb. We avoid it with a symmetric reconstruction: sign-align the halves of each mode 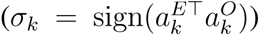, form one shared coefficient 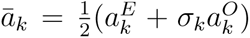, and place both halves of the mode in a single full-band signature *v_k_* with 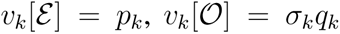, reconstructed against a single grand mean 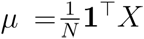.

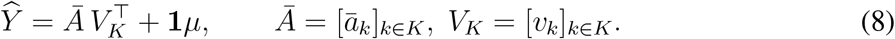

When the two views carry independent noise of equal variance, averaging the two coefficient estimates additionally halves their noise variance, a secondary denoising gain on top of comb removal. The resulting cube enters the shared post-processing (Section 3.5) on the same footing as every other method.

### 3.5 Post-processing and evaluation

Every denoised dispersive cube is carried through the same post-processing chain. We first denormalize the cube back to physical Re{*χ*} units, then retrieve the Raman spectrum Im{*χ*} with a Hilbert transform^48^ followed by an asymmetric-least-squares phase-error correction that removes the residual slow phase distortion left by an imperfect NRB estimate.^11^ The corrected Raman spectrum is the quantity from which the biological Raman signal is read and on which the denoisers are evaluated.

We evaluate all denoisers using a common protocol within each dataset. On the simulated cubes, whose complex ground truth is known exactly, we report peak signal-to-noise ratio (PSNR) and structural similarity index (SSIM),^49^ both evaluated on the retrieved resonant lineshape. PSNR and SSIM are each reported three times: over the whole image, and separately over a foreground and a background region. The foreground mask is built from the ground-truth resonant intensity by retaining the largest connected component of the thresholded map and taking its convex hull, which yields a compact, hole-free region enclosing the sample; the background is its complement. The split matters because the background carries no resonant signal: a whole-image score averages over pixels where there is nothing to recover, so a gain realized there reflects the flattening of empty space but not the restoration of a band. We therefore quote the whole-image values as the summary metric and use the foreground–background decomposition diagnostically, to establish where each method’s improvement is realized. The foreground is the region that carries the chemical information, and a method whose gain is concentrated in the background has improved the image without improving the measurement. For glycerol, we report the PSNR against a common, galvo-resolved pseudo-ground truth constructed directly from averaging noisy measurements. After VST and Whittaker detrending, we average 100 slow-axis rows, producing one reference spectrum at each galvo position. The averaged spectra undergo the same post-processing, and each output is matched to the reference by a global scale and offset correction before the PSNR is computed over the full crop. This reference shares measurement noise with the input and is treated as an approximate reference rather and is not an independent ground truth. For this dataset, structural similarity is omitted because averaging removes spatial structure from the reference. For images of polystyrene beads, we fit the full width at half maximum (FWHM) of a bead profile to compare each denoiser’s apparent profile width with that of the noisy input. For experimental *C. elegans*, no ground truth is available. We therefore compare the reconstructed Raman maps qualitatively, displaying corresponding regions side by side with a common intensity scale. We examine visible background fluctuations and the appearance of fine spatial features, characterizing differences between the outputs without assigning a quantitative fidelity ranking.

## 4 Results

### 4.1 Simulated BCARS with ground truth

The simulated BCARS cubes allow for a direct comparison of the denoised spectra against the true ground truth (GT) on the resonant (Im{*χ*_R_}) lineshape. We evaluate the performance of each denoising method using PSNR and SSIM, each computed over the whole image and separately over the foreground and background regions (Section 3.5). Fig. 3 shows the magnitude and spatial distribution of mean squared error (MSE) over the image foreground: for every denoiser, the upper panel maps the per-pixel error against the GT and the lower panel shows the difference between the mean spectral power of the denoised and GT spectra, with excess power (blue) and deficit (red) indicating where a method adds or removes spectral structure relative to the GT. The methods with the lowest foreground MSE (DDS2M, DIP, and CCV) show smaller power differences at mid-range spectral frequencies, and DDS2M also at the higher frequencies. The corresponding reconstructed images and single-pixel spectra are shown in Fig. S2 (SI).

**Fig 3.**
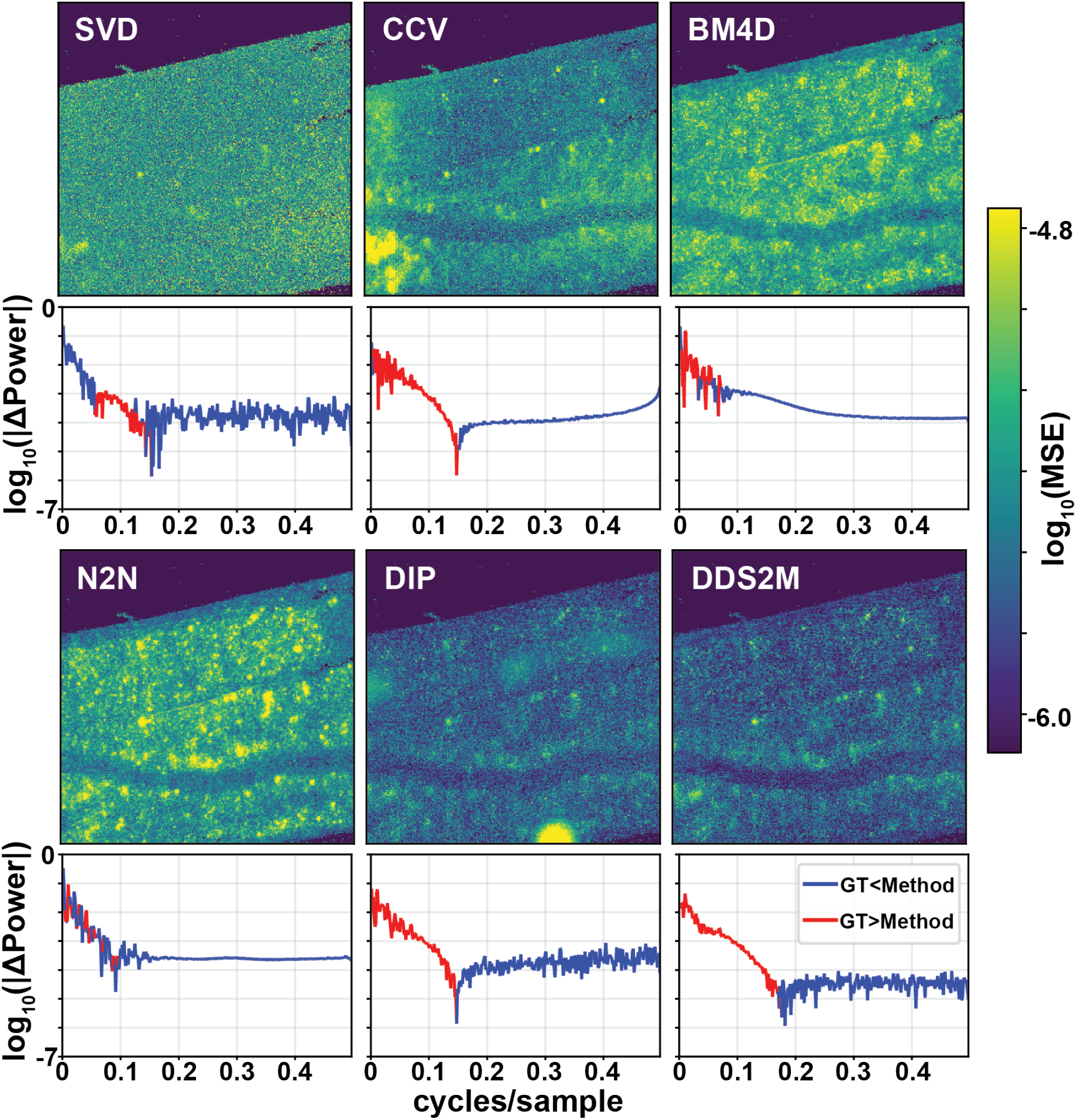
Error structure of six denoisers on the simulated cube. For each method, the upper panel maps per-pixel log_10_(MSE) against the ground truth; the lower panel plots the corresponding difference in mean spectral power, log_10_ |ΔPower|. Blue indicates excess power relative to GT while red indicates a deficit.

The results are summarized in Table 1. On whole-image PSNR, each denoiser is at least 8.5 dB above the noisy input. The ranking resolves roughly into three tiers: DDS2M (41.43 dB) leads the next-ranked method by 1.5 dB; DIP, N2N, BM4D, and CCV (39.96, 39.62, 39.46, 39.20 dB) lie within 0.8 dB of one another, less than the spread across cubes; and SVD (37.86 dB) trails CCV by 1.3 dB.

**Table 1.** Six denoisers and the noisy input on the simulated cubes, scored on Raman (Im *χ*_R_) lineshapes against the exact ground truth. Metrics are reported over the whole image and separately over the foreground (signal-bearing) and background regions. Values are mean ± standard deviation across five simulated data cubes.

| Method | Whole image |  | Foreground |  | Background |  |
| --- | --- | --- | --- | --- | --- | --- |
|  | PSNR (dB) | SSIM | PSNR (dB) | SSIM | PSNR (dB) | SSIM |
| DDS2M | $41.43 \pm 0.69$ | $0.475 \pm 0.041$ | $39.90 \pm 0.85$ | $0.523 \pm 0.060$ | $42.40 \pm 1.15$ | $0.434 \pm 0.034$ |
| DIP | $39.96 \pm 0.83$ | $0.488 \pm 0.056$ | $40.14 \pm 0.76$ | $0.556 \pm 0.064$ | $39.67 \pm 1.16$ | $0.440 \pm 0.038$ |
| N2N | $39.62 \pm 0.93$ | $0.382 \pm 0.034$ | $37.69 \pm 0.40$ | $0.393 \pm 0.045$ | $41.32 \pm 1.75$ | $0.373 \pm 0.024$ |
| BM4D | $39.46 \pm 1.09$ | $0.345 \pm 0.035$ | $37.78 \pm 0.70$ | $0.353 \pm 0.049$ | $40.81 \pm 1.71$ | $0.337 \pm 0.022$ |
| CCV | $39.20 \pm 1.00$ | $0.455 \pm 0.056$ | $38.78 \pm 1.44$ | $0.556 \pm 0.053$ | $39.22 \pm 1.10$ | $0.382 \pm 0.046$ |
| SVD | $37.86 \pm 1.40$ | $0.396 \pm 0.078$ | $37.65 \pm 1.53$ | $0.483 \pm 0.079$ | $37.86 \pm 1.45$ | $0.339 \pm 0.047$ |
| Noisy | $29.30 \pm 1.21$ | $0.127 \pm 0.024$ | $26.61 \pm 0.58$ | $0.110 \pm 0.010$ | $32.24 \pm 3.16$ | $0.133 \pm 0.029$ |

Gains per method relative to the noisy input are reported in Fig. 6.

Whole-image SSIM produces a slightly different tier structure with DIP, DDS2M, and CCV (0.488, 0.475, and 0.455, respectively) within 0.035 of one another. SVD falls at an intermediate value (0.396) with N2N and BM4D (0.382, 0.345) trailing.

These whole-image numbers conceal where each method’s gain is realized. Comparing fore-ground and background performance reveals that DIP, CCV, and SVD focus relatively more on larger foreground signals than on the background compared to the other methods. In foreground PSNR, DDS2M and DIP perform equally with CCV slightly behind, and SVD, BM4D and N2N tightly grouped together and behind CCV. In background rankings, DIP, CCV, and SVD all fall behind DDS2M, BM4D and N2N. Changes in SSIM foreground and background performance are similar except that DIP retains a leading SSIM score for background. We note that, while the backgrounds in these images contain no information of interest, they can be seen as surrogates for small signals in an image with very high dynamic range, so we can anticipate that DDS2M, BM4D and N2N may recover those small signals better than the other three methods.

### 4.2 Glycerol: comparison against a galvo-resolved pseudo-GT

The homogeneous glycerol sample provides a measurement-based pseudo-ground-truth constructed by averaging slow-axis rows while retaining galvo position (Section 3.5). Fig. S4 shows representative single-pixel Raman spectra, with the reference taken at the corresponding galvo position. The annotated PSNR values are evaluated against this common reference.

BM4D achieves the highest PSNR at 32.97 dB, followed by DDS2M (31.55), DIP (30.76), SVD (30.16), CCV (29.97), and N2N (28.21), compared with 20.46 dB for the noisy input. The ordering differs from the simulated tissue benchmark. These results describe agreement with the averaged glycerol reference but not establishing a general ordering of reconstruction fidelity across samples.

### 4.3 Polystyrene beads: spatial fidelity and artifacts

The polystyrene beads provide a test of spatial fidelity and artifact generation in the absence of an independently measured ground truth, but their sharp spatial features allow for a visual assessment of the effect of denoising on the spatial fidelity. Fig. 4 shows the noisy input and the denoised images at the 1004 cm*^−^*^1^ band, with the boxed bead and its *X*/*Y* profile axes marked. The top right panels show the boxed bead after each denoiser, annotated with the fitted FWHM; the bottom panels show the normalized intensity profiles through the bead center along *X* and *Y*, with Gaussian fits.

**Fig 4.**
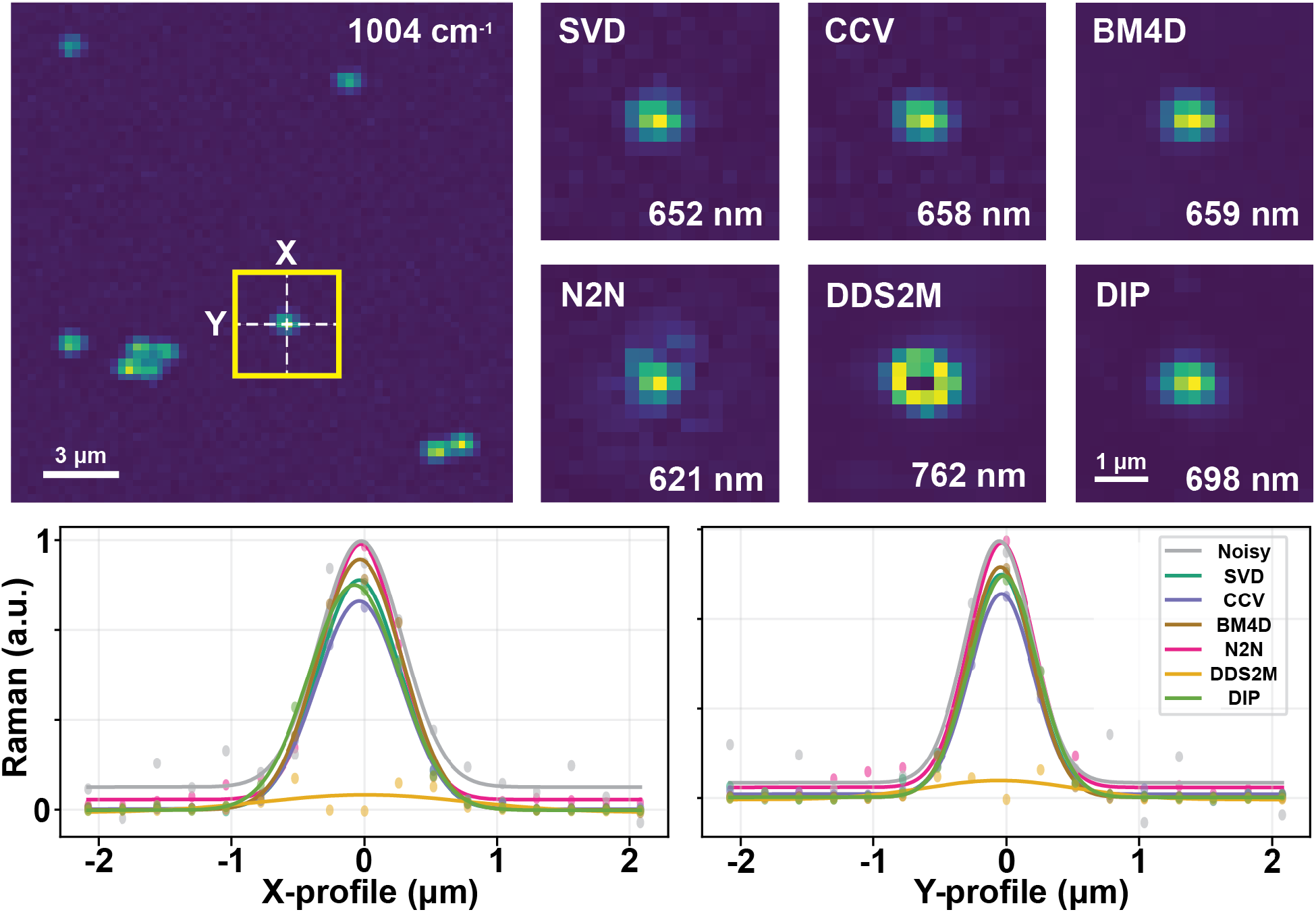
Spatial fidelity on polystyrene beads. Top left: resonant image at the 1004 cm*^−^*^1^ polystyrene band (noisy), with the analyzed bead boxed and its *X*/*Y* profile axes marked; 3 *µ*m scale bar. Top right: the boxed bead after each denoiser, annotated with the fitted FWHM; 1 *µ*m scale bar. Bottom: normalized intensity profiles through the bead center along *X* and *Y*, with Gaussian fits. SVD, CCV, and BM4D preserve the bead profile (FWHM within ∼10 nm of the 658 nm noisy estimate). DIP broadens modestly (698 nm), and DDS2M both broadens the profile (762 nm) and suppresses peak amplitude, producing an artifact consistent with its clipped, bounded range at this strong band. N2N distorts the profile: with the bead cube in its training set a single-Gaussian fit reads 621 nm, narrowed by a negative ring undershoot; with it excluded, a positive overshoot broadens the fit to 1075 nm (Fig. S5). In neither case does the fitted width reflect a faithfully recovered profile.

SVD, CCV, and BM4D yield fitted FWHM values within approximately 10 nm of the 658 nm estimate from the noisy input. DIP yields a wider fitted profile of 698 nm. N2N distorts the profile in opposite directions depending on whether the bead cube was or was not included in the training set. With the bead cube included, it returns a narrower profile than any other method, 621 nm, an apparent sharpening produced by a negative ring undershoot at the bead edge rather than by recovered resolution; with the bead cube excluded it broadens and distorts the profile to 1075 nm (Fig. S5). DDS2M suppresses the peak amplitude and returns a broadened artifact (FWHM 762 nm), consistent with the clipping and bounded output used in our DDS2M implementation (see Sec. 5).

### 4.4 C. elegans: biological application

The experimental *C. elegans* cube provides heterogeneous tissue without an independently measured ground truth. Fig. 5 compares the noisy Raman map with the six denoised reconstructions at the 1655 cm*^−^*^1^ C=C stretching band. All methods reduce visible background fluctuations and make the gross tissue organization easier to distinguish. The reconstructions differ in local contrast and texture, but without an independently measured ground truth, these differences do not establish a ranking of reconstruction fidelity.

**Fig 5.**
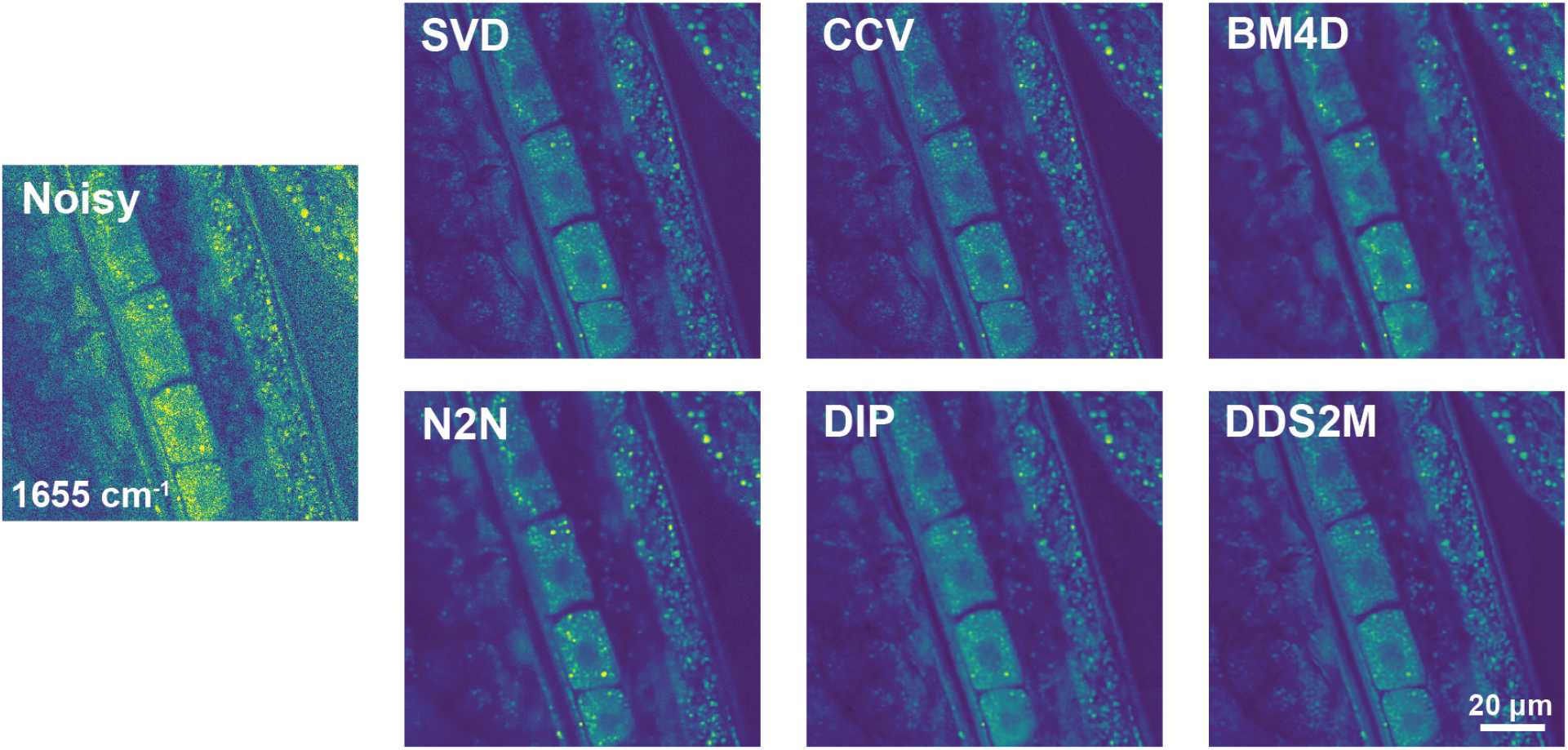
Biological application to *C. elegans* tissue. Resonant Raman maps of a heterogeneous *C. elegans* region for the noisy input and all six denoisers at 1655 cm*^−^*^1^; 20 *µ*m scale bar.

## 5 Discussion

### 5.1 A practical trade-off map

Fig. 6 recasts the fidelity of Table 1 against cost and operator burden. Each panel plots a foreground fidelity gain over the noisy input against wall-clock runtime, for **(A)** PSNR and **(B)** SSIM, with marker shape and color encoding operator burden. SSIM gains are taken on a logit scale, log_10_[SSIM*/*(1 − SSIM)], because all observed SSIM values lie in (0, 1), where differences near the upper end are compressed; both transforms are defined in the Supplement. The corresponding background gains are shown in Fig. S3 (SI).

**Fig 6.**
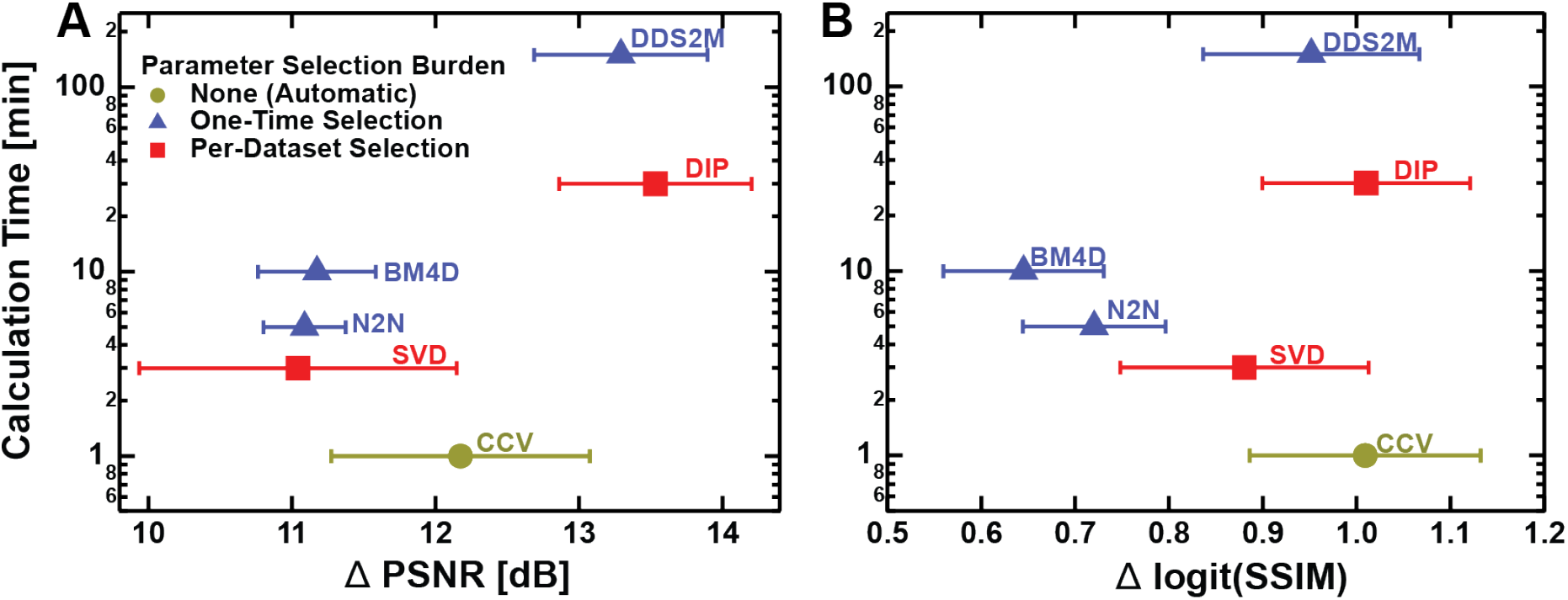
Fidelity–cost–burden trade-off on the simulated cubes. *Improvement over the noisy input*, evaluated in the foreground region, is plotted against wall-clock runtime for **(A)** PSNR, evaluating signal fidelity via pixel-level reconstruction accuracy, and **(B)** SSIM, measuring spatial–spectral fidelity, on a logit scale Δ log_10_[SSIM*/*(1 − SSIM)]. Both horizontal axes are differences against the noisy input rather than absolute scores, so a larger value denotes a larger gain from denoising, not a higher absolute fidelity. Marker shape and color encode operator burden, defined by how often a human must make a subjective, cube-specific choice: automatic (no per-cube tuning), one-time (a parameter or training run fixed once and reused unchanged), and per-dataset (a choice remade for every cube). The error bars represent the standard deviation across five simulated data cubes. The corresponding gains in the background region are shown in Fig. S3 (SI).

The two axes are complementary. PSNR quantifies point-wise fidelity, penalizing all deviations equally regardless of their spatial or spectral context; it is sensitive to global intensity offsets and broadband noise variance but agnostic to the spatial arrangement of the errors. SSIM instead compares local means, variances, and covariances within a sliding window, and so specifically penalizes the loss of local structure such as edges and texture. A method that over-smooths spatial features can therefore hold its PSNR while its SSIM degrades.

Figure 6 reports foreground gains to focus on the signal-bearing region; Fig. S3 reports the corresponding background gains. All methods achieve a larger PSNR gain in the foreground than in the background. The difference is 5.2–6.1 dB for DIP, SVD, and CCV, compared with 2.0–3.1 dB for DDS2M, BM4D, and N2N.

DIP leads on both fidelity axes and DDS2M on PSNR, but at 30 and 150 min they run one to two orders of magnitude slower than the rank-domain methods, and DIP requires an early-stopping decision remade for each cube. CCV runs in about a minute and ranks third in mean foreground PSNR gain. Its mean logit-SSIM gain is close to that of DIP (1.009 ± 0.123 versus 1.010 ± 0.111). It selects components automatically using a fixed correlation threshold, without per-cube parameter adjustment. SVD takes the next rank position with respect to fidelity and time, but needs a per-cube rank choice. N2N is the only method that must be trained, which is not shown by its one-time runtime marker, and has a spatial fidelity that depends on the training distribution (Sec. 4.3). Finally, DDS2M’s bounded output range suppresses sparse, high-amplitude bands, a failure that aggregate metrics on a single cube do not register (Sec. 4.3).

### 5.2 Failure modes and costs

CCV is fast, but, like our implementation of N2N, the method rests on the assumption that adjacent spectral channels carry independent noise (Sec. 3.4). If that assumption were to fail, for example in the presence of spectrally correlated noise, the correlated noise would reproduce across the odd/even split and would be misinterpreted as signal, leading to an overestimation of the rank and a corresponding under-denoising. It may be possible to ameliorate this with appropriate post-hoc Gaussian filtering. The apparently elevated noise power at high frequency in the CCV panel of Fig. 3 may represent a case of unaddressed spectrally correlated noise.

BM4D ranks fourth in whole-image PSNR and last in whole-image SSIM. Its foreground SSIM exceeds its background SSIM by 0.016, the smallest difference among the six methods. One possible explanation is that collaborative filtering suppresses local image variation along with noise. We did not vary the shrinkage threshold to test this explanation. BM4D also achieves the highest PSNR against the averaged glycerol reference (Section 4.2), where spatial averaging is well suited to the homogeneous sample.

DDS2M suppresses the strong polystyrene band (Fig. 4). DDS2M rescales its input to [−1, 1] and clamps its reconstruction to [0, 1], so normalized amplitudes above the upper percentile cannot be recovered; we therefore run it on the clipped cube (Section 3.2), which removes the top of sparse, high-amplitude peaks. On the unclipped cube, the sparse peak instead sets the normalization scale, and the reconstruction retained mainly the brightest pixels. The suppression therefore reflects the bounded range of this implementation rather than an established property of diffusion denoising.

N2N shows the same pattern as BM4D, more strongly: it scores 3.63 dB higher in the background than in the foreground, the largest gap of the six, and its foreground SSIM advantage over the background is only 0.020. The bead comparison additionally reveals sensitivity to the training data. Including the bead cube in training produces a negative ring undershoot and a fitted FWHM of 621 nm. Excluding it produces a distorted, broader profile with a fitted FWHM of 1075 nm (Fig. S5). These observations show that the training composition affects the reconstructed profile, while including the bead data does not eliminate artifacts. N2N also has the lowest PSNR against the glycerol reference (Section 4.2).

DIP achieves high foreground fidelity at a suitable stopping iteration. Too few iterations underfit the signal, whereas continued fitting can reproduce noise. Here, the stopping iteration is selected by visual inspection for each cube. This requirement adds operator input to the reconstruction. Ground-truth-free stopping remains a challenge for DIP methods;^31, 32^ the suitability of proposed criteria^50^ for BCARS remains to be evaluated.

### 5.3 Limitations

Three limitations bound the scope of these conclusions.

First, the simulated benchmark contains five cubes, with metrics reported as means and standard deviations across cubes. Each experimental sample is represented by a single acquisition. Experimental variability is therefore not quantified. The glycerol PSNR and bead FWHM values have no uncertainty estimates, and the *C. elegans* comparison is qualitative.

Second, all experimental cubes were acquired on one instrument and cover only the fingerprint region. The same system also records the CH-stretch region, so the restriction here is a scoping decision, not an instrumental one. The fingerprint is where denoising is most consequential, because weak, densely overlapping resonances sit on a comparatively large nonresonant background and per-channel SNR is correspondingly low. It is also where the front-end is on safest ground, since the linearization underlying the transform assumes |*χ*_R_| ≪ *χ*_NR_ (Section 2), and Fig. 2 confirms that the stabilized variance is flat across the fingerprint, its only departure lying at the Raman-silent edge and traceable to read noise than to the neglected quadratic term. Extending the benchmark to the CH-stretch region is therefore not a matter of running the same pipeline on additional cubes: at the strongest lipid bands |*χ*_R_| is no longer small compared with *χ*_NR_, so the stabilization would first have to be re-validated in a regime where the linearization is not guaranteed. The accuracy of variance stabilization and the denoiser rankings in this regime remain to be evaluated. More generally, the transform follows from the BCARS forward model and is not tied to a particular setup, but the numerical rankings reported here reflect the instrument’s excitation profile, and to a much lesser extent the detector’s (relatively small) read noise and, in principle, relative fidelity rankings may vary slightly on systems with different background envelopes.

Third, the benchmark compares denoisers on the whitened cube without isolating the contribution of whitening itself. Every method is run behind the same variance-stabilizing front-end, which is what allows the measured differences to be attributed to the algorithms rather than to their differing tolerance of heteroscedastic noise; it does not establish that an unstabilized comparison would have ordered the methods differently. Quantifying that would require rerunning each denoiser on the raw cube, which we leave to future work. For the same reason we do not claim that the transform improves any individual denoiser, only that it supplies a common statistical footing on which they can be compared.

## 6 Conclusion

We developed a BCARS denoising benchmark with a physics-derived variance-stabilizing transform and a shared phase-retrieval pipeline. We also introduced CCV, which uses cross-covariance between redundant spectral views to suppress noise and automatically select components. In simulations, the transform reduced the ratio of maximum to minimum channel noise variance from approximately 72 to 1.2.

Performance depended on the sample, metric, and image region. DDS2M and DIP achieved the highest mean foreground PSNR on the simulated cubes, whereas CCV matched DIP’s mean foreground SSIM with substantially shorter runtime. Experimental comparisons identified sharp-feature distortions and a different ranking against the averaged glycerol reference. These results support evaluating denoisers using both reference-based metrics and sample-specific tests.

CCV provides a practical option when runtime and automatic operation are priorities. DIP requires a stopping decision for each cube, while the bounded range of the DDS2M implementation used here limits its recovery of sparse, high-amplitude bands. These conclusions apply to the fingerprint-region datasets and implementations evaluated in this study.

The testing framework generalizes beyond the six methods tested here. Because the variance stabilized cube already carries physical Re *χ*_R_ units, the causality constraint currently imposed after restoration could instead be folded into it, so that Kramers–Kronig consistency regularizes the denoiser rather than correcting its output. RamPINN^51^ has demonstrated this principle for the adjacent problem of separating the resonant signal from the nonresonant background in CARS, enforcing the Kramers–Kronig relations through a differentiable Hilbert-transform loss; applying the same construction to restoration of variance-stabilized BCARS data could extend this approach. The split-half reproducibility that sets the rank in CCV is likewise not specific to a low-rank model, and the same ground-truth-free reliability statistic may offer a principled stopping rule for the untrained priors whose optimum is presently found by eye. Finally, we have evaluated the denoisers only for the fingerprint region. Extending the front-end and the benchmark to the CH-stretch region, where the linearization must first be re-validated, is the natural next step.

## Code, Data, and Materials Availability

The datasets supporting this study are openly available on Zenodo at DOI 10.5281/zenodo.22968812. The code is available on GitHub at https://github.com/CiceroneLab/bcars-vst-denoise.

## Acknowledgments

This work was funded by the Department of Energy Biological and Environmental Research (DOE BER DE-SC0022121).

