## Supplemental Material for "Denoising broadband CARS hyperspectral images on variance-stabilized data: a unified benchmark"

**Haoyu Xu[a], Marcus T. Cicerone[a,b,*]**
[a]The Wallace H. Coulter Department of Biomedical Engineering, Georgia Institute of Technology, Atlanta, GA 30332, USA
[b]School of Chemistry & Biochemistry, Georgia Institute of Technology, Atlanta, GA 30332, USA

*Marcus T. Cicerone,

### Supplementary information

#### *Metric definitions*

All simulated-cube metrics are computed on the retrieved resonant lineshape $\mathrm{Im}\,\chi_{\mathrm{R}}$ against the exact ground truth, and are reported over the whole image and separately over the foreground and background regions defined in Section 3.5.

Peak signal-to-noise ratio is

$$\mathrm{PSNR} = 10\log_{10}\left(\frac{L^2}{\mathrm{MSE}}\right), \qquad \mathrm{MSE} = \frac{1}{N}\sum_{i=1}^{N}(x_i - y_i)^2, \tag{S1}$$

where $x_i$ and $y_i$ are corresponding values in the ground-truth and denoised cubes, $N$ is the number of voxels in the region being scored, and $L$ is the dynamic range of the ground truth. PSNR quantifies point-wise fidelity: it penalizes all deviations equally, regardless of their spatial or spectral context.

The structural similarity index instead compares local statistics within a window:[1]

$$\mathrm{SSIM}(x,y) = \frac{(2\mu_x\mu_y + C_1)(2\sigma_{xy} + C_2)}{(\mu_x^2 + \mu_y^2 + C_1)(\sigma_x^2 + \sigma_y^2 + C_2)}, \tag{S2}$$

where $\mu_x$ and $\mu_y$ are local means, $\sigma_x^2$ and $\sigma_y^2$ local variances, $\sigma_{xy}$ the local covariance, and $C_1 = (K_1 L)^2$, $C_2 = (K_2 L)^2$ are stabilization constants with the default parameters. The scalar SSIM is obtained by averaging the local values over all spatial positions; for a cube of shape $(n_y, n_x, n_\nu)$ we compute SSIM independently at each spectral channel and average over the $n_\nu$ channels.

Fig. 6 reports gains relative to the noisy input rather than absolute scores. For PSNR the gain is the plain difference, $\Delta\mathrm{PSNR} = \mathrm{PSNR}_{\mathrm{method}} - \mathrm{PSNR}_{\mathrm{noisy}}$, with both terms evaluated over the same region. All observed SSIM values lie in $(0, 1)$, where differences near the upper end are compressed, so the gain is taken on a logit scale,

$$\Delta\mathrm{logit}(\mathrm{SSIM}) = \log_{10}\frac{\mathrm{SSIM}_{\mathrm{method}}}{1-\mathrm{SSIM}_{\mathrm{method}}} - \log_{10}\frac{\mathrm{SSIM}_{\mathrm{noisy}}}{1-\mathrm{SSIM}_{\mathrm{noisy}}}, \tag{S3}$$

which makes equal improvements comparable across regions whose noisy baselines differ.

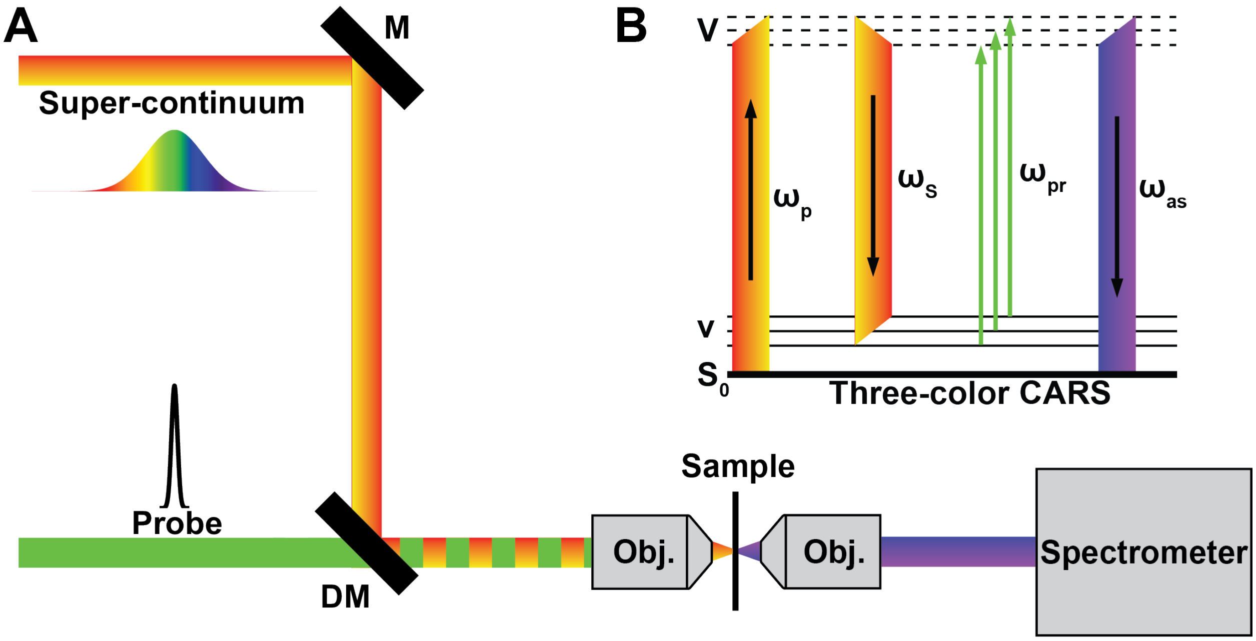


**Fig S1 Simplified BCARS setup and three-color imaging.** Schematic of the broadband CARS microscope: a narrowband probe and a supercontinuum Stokes beam, generated in a photonic-crystal fiber, are combined to impulsively excite the vibrational bands of the fingerprint region (400–1800 $cm^{-1}$) and generate the anti-Stokes signal dispersed onto the spectrometer. The three-color excitation places the excited band across the fingerprint window, so that a full fingerprint spectrum is recorded at every pixel in a single acquisition.

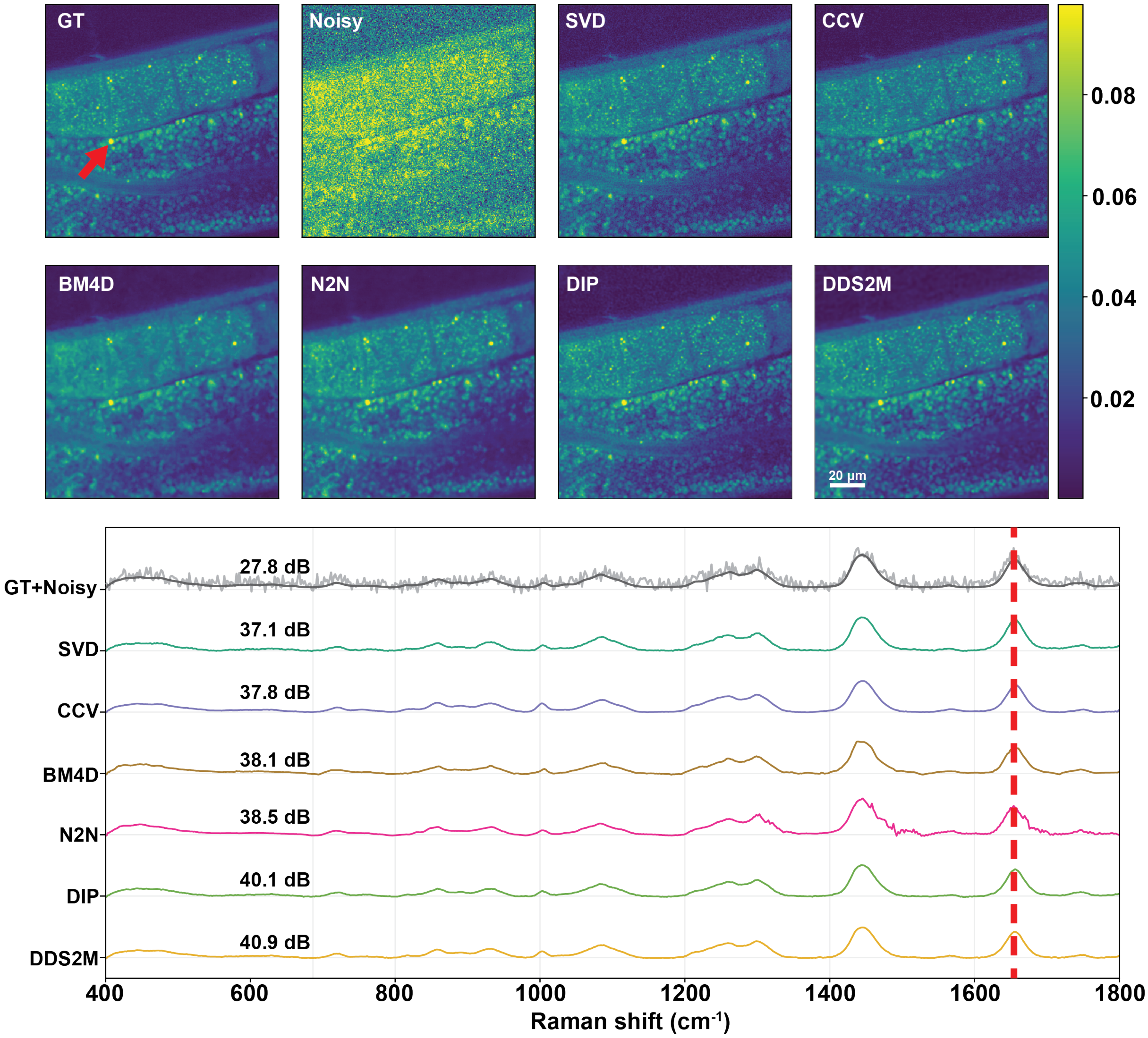


**Fig S2 Benchmark on the simulated cube with exact ground truth.** Top: resonant ($\mathrm{Im}\,\chi_{\mathrm{R}}$) images at a representative Raman band for the complex ground truth (GT), the noisy input, and all six denoisers, on a shared color scale (right); 20 $\mu$m scale bar. Bottom: single-pixel resonant spectra at the pixel arrowed in the GT panel, offset vertically for clarity and each annotated with its resonant-domain PSNR (dB) against the known ground truth (Table 1); the noisy trace (light grey) is overlaid on the GT (dark). DDS2M and DIP lead, followed by N2N, BM4D, CCV, and SVD, all well above the noisy input. The dashed red line marks the band used for the images.

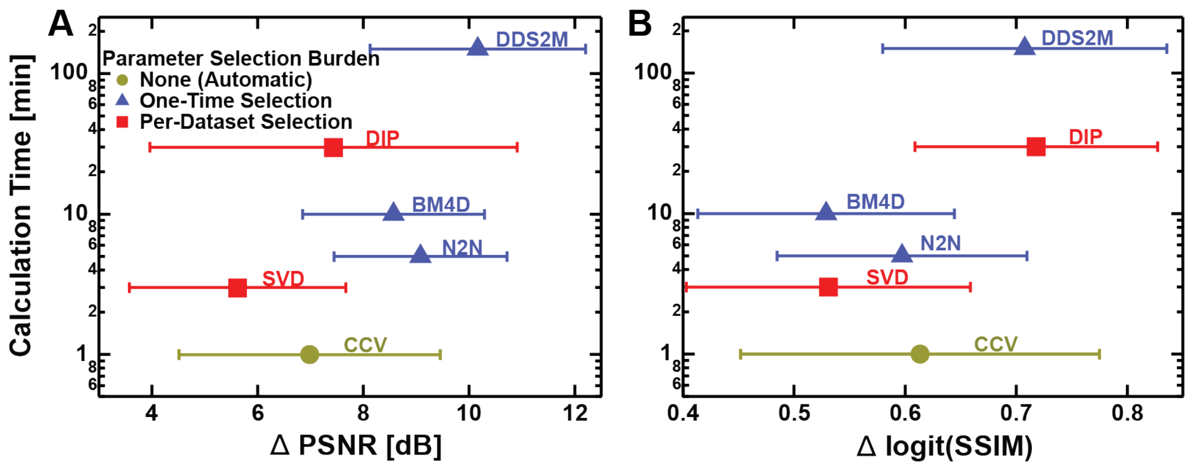


**Fig S3 Fidelity–cost–burden trade-off in the background region.** As Fig. 6, but with the improvement over the noisy input evaluated over the background rather than the foreground; axes, markers, and error bars are defined as in that figure.

**Table S1** Configuration of the three deep-learning denoisers on the simulated benchmark (Table 1): model, key hyperparameters, and any modification from the reference implementation.

| Method | Model | Key hyperparameters | Modification from reference |
|---|---|---|---|
| **DDS2M** [2] | Self-supervised diffusion (DDRM-style) with an untrained spatio-spectral prior: each rank component pairs a spatial U-Net with a spectral fully-connected network. | rank 10; $\eta = 1.0$; 2000 timesteps (linear $\beta$ from $1 \times 10^{-4}$ to $5 \times 10^{-3}$); learning rate $5 \times 10^{-4}$; spatial TV disabled. | Applied to the VST-whitened cube; the output is clamped to $[0, 1]$, so peak amplitudes are limited on sparse-peak cubes. |
| **DIP** [3] | Untrained encoder–decoder network (depth 4, 128 channels) fit per cube; total-variation regularization along the wavenumber axis. | spectral-TV weight $5 \times 10^{-3}$ (order 1); learning rate $5 \times 10^{-3}$; best early-stop checkpoint. | Adds a 1-D spectral-TV term along the band axis to the reference (spatial-only) prior; early stopping is the primary lever. |
| **N2N** [4] | 3-D convolutional U-Net (depth 4, 32 base filters, $3 \times 3 \times 3$ kernels, residual output). | MAE loss; learning rate $4 \times 10^{-4}$ with ReduceLROnPlateau (factor 0.5, patience 10); batch size 16; 100 epochs $\times$ 100 steps; patches $32 \times 32 \times 96$, stride $16 \times 16 \times 48$, augmented with 90° rotations and flips. | Trained self-supervised on odd/even spectral splits of the VST-whitened cube (two independent noisy views of the same scene). |

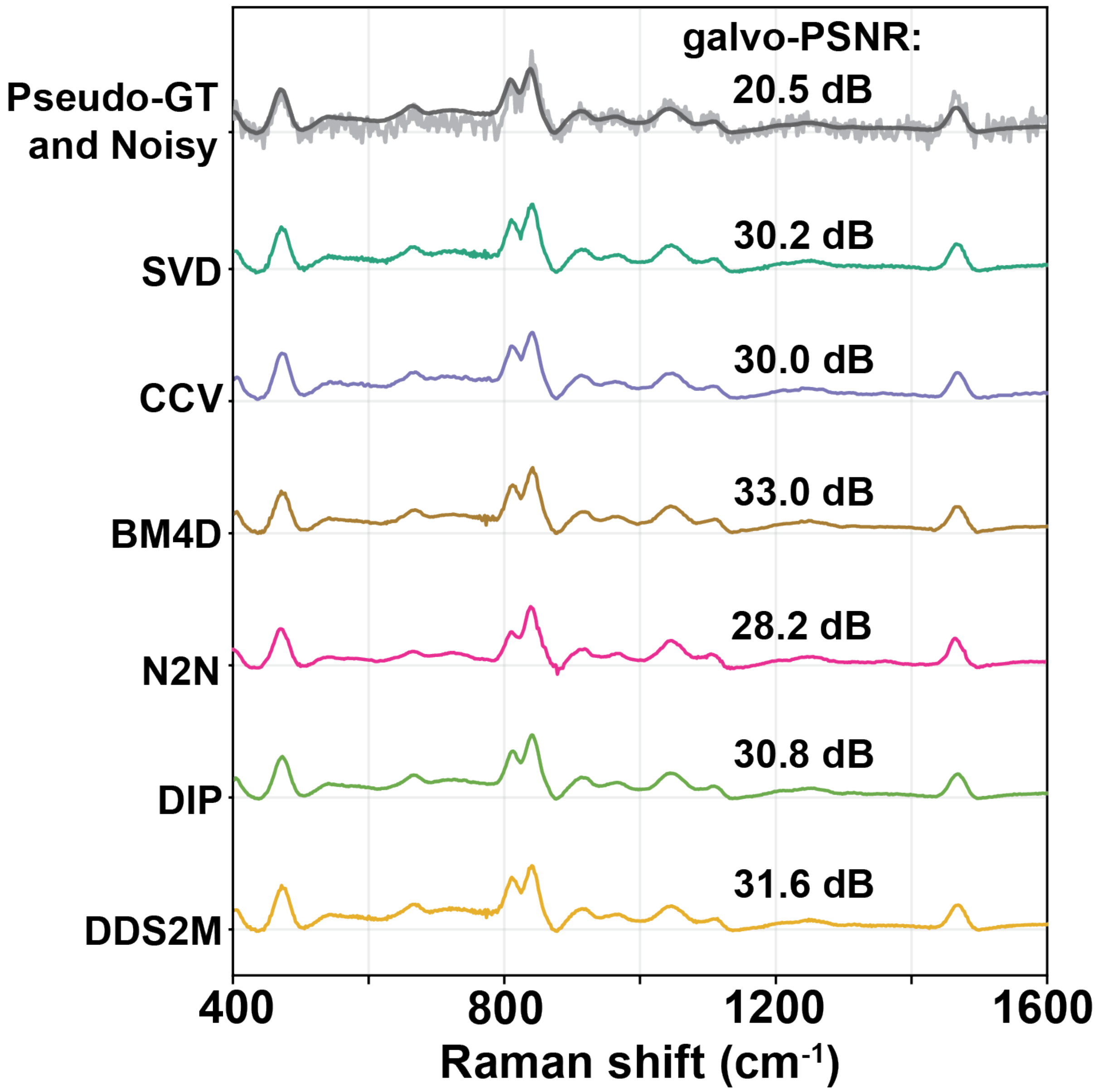


**Fig S4 Denoising on the homogeneous glycerol cube.** Representative Raman spectra for the six denoisers, offset vertically, each annotated with its PSNR against the noisy average at the corresponding galvo position. The top trace overlays the noisy spectrum (light) on the reference (dark).

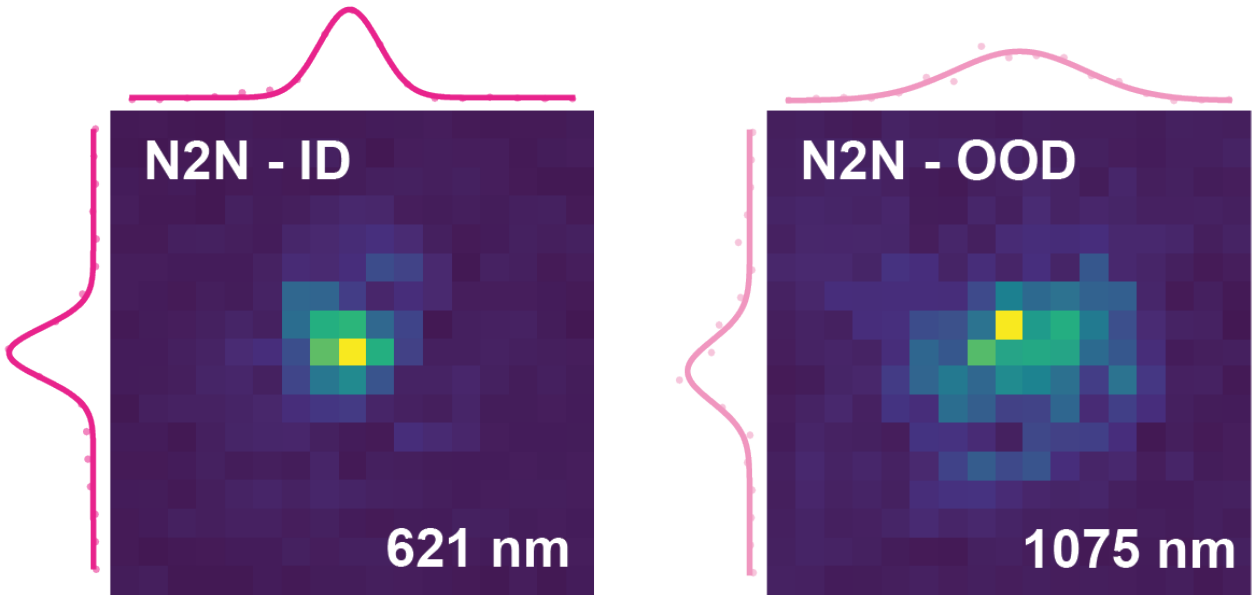


**Fig S5 N2N failure mode in the beads data.** Left: bead cube included in the training set; the fitted FWHM is 621 nm, narrowed by a negative ring undershoot. Right: bead cube excluded from the training set; the fitted FWHM is 1075 nm, broadened by a positive ring overshoot. Both fitted widths are artifacts of the ringing, not faithful recoveries of the bead profile, illustrating N2N's sensitivity to features absent from its training distribution.

*References*

1 Zhou Wang, A.C. Bovik, H.R. Sheikh ... E.P. Simoncelli, “Image quality assessment: from error visibility to structural similarity,” *IEEE Transactions on Image Processing* **13**(4), 600–612 (2004).

2 Yuchun Miao, Lefei Zhang, Liangpei Zhang ... Dacheng Tao, “DDS2M: Self-supervised denoising diffusion spatio-spectral model for hyperspectral image restoration,” in *2023 IEEE/CVF International Conference on Computer Vision (ICCV)*, 12052–12062, IEEE (2023).

3 Dmitry Ulyanov, Andrea Vedaldi, and Victor Lempitsky, “Deep image prior,” in *Proceedings of the IEEE Conference on Computer Vision and Pattern Recognition (CVPR)*, 9446–9454 (2018).

4 Jaakko Lehtinen, Jacob Munkberg, Jon Hasselgren ... Timo Aila, “Noise2Noise: Learning image restoration without clean data,” in *Proceedings of the 35th International Conference on Machine Learning (ICML)*, *PMLR* **80**, 2965–2974 (2018).
